# Training for object recognition with increasing spatial frequency: A comparison of deep learning with human vision

**DOI:** 10.1101/2021.01.24.427905

**Authors:** Lev Kiar Avberšek, Astrid Zeman, Hans P. Op de Beeck

## Abstract

The ontogenetic development of human vision, and the real-time neural processing of visual input, both exhibit a striking similarity – a sensitivity towards spatial frequencies that progress in a coarse-to-fine manner. During early human development, sensitivity for higher spatial frequencies increases with age. In adulthood, when humans receive new visual input, low spatial frequencies are typically processed first before subsequently guiding the processing of higher spatial frequencies. We investigated to what extent this coarse-to-fine progression might impact visual representations in artificial vision and compared this to adult human representations. We simulated the coarse-to-fine progression of image processing in deep convolutional neural networks (CNNs) by gradually increasing spatial frequency information during training. We compared CNN performance, after standard and coarse-to-fine training, with a wide range of datasets from behavioural and neuroimaging experiments. In contrast to humans, CNNs that are trained using the standard protocol are very insensitive to low spatial frequency information, showing very poor performance in being able to classify such object images. By training CNNs using our coarse-to-fine method, we improved the classification accuracy of CNNs from 0% to 32% on low-pass filtered images taken from the ImageNet dataset. When comparing differently trained networks on images containing full spatial frequency information, we saw no representational differences. Overall, this integration of computational, neural, and behavioural findings shows the relevance of the exposure to and processing of input with a variation in spatial frequency content for some aspects of high-level object representations.

## Introduction

The role of spatial frequency has been extensively researched in the development of human vision, with much effort being directed towards visual acuity and contrast sensitivity (Leat, et al., 2009; Norcia & Tyler, 1985; Norcia, Tyler & Hammer, 1990, Banks & Salapatek, 1978; Benedek et al., 2003; Ellemberg et al., 1998; Mayer & Dobson, 1982; Peterzell et al., 1995; Stiers et al., 2003). Visual acuity can be classified into recognition (perceived detail) and resolution (the separation between dots or gratings i.e. spatial frequency) that a person can successfully resolve. Contrast sensitivity is the smallest difference in luminance that can be perceived between an object and its immediate surroundings. Contrast sensitivity is measurable across the whole spectrum of spatial frequencies, referred to as the contrast sensitivity function or CSF (Leat et al., 2009). Researchers have used a variety of methods, including visual evoked potential (VEP) (Norcia & Tyler, 1985; Norcia, Tyler & Hammer, 1990) and psychophysiological methods (Banks & Salapatek, 1978; Benedek et al., 2003; Ellemberg et al., 1998; Mayer & Dobson, 1982; Peterzell et al., 1995; Stiers et al., 2003), to reach similar conclusions: that visual acuity and contrast sensitivity increase with age. At an infant level, visual acuity and contrast sensitivity are very poor, and over the span of a number of years, they improve to reach a high functioning level, usually seven to twelve years for visual acuity and eight to nineteen years for contrast sensitivity (according to Leat et al., 2009). The peak of infants’ contrast sensitivity, as well as the point at which their contrast perception falls to zero (their cut-off frequency), are both lower than that of adults (Kiorpes, 2016; Leat et al., 2009).

In addition to this coarse-to-fine progression during the ontogenetic development of human vision, there appears to be a similar tendency in the real-time neural processing of fully mature adults as their visual system receives input moment-to-moment. Modern theories of vision suggest that the visual system works in a coarse-to-fine manner, in which the low spatial frequency content (LSF) of visual input, which contains coarse information such as global shape, takes precedence over high spatial frequency (HSF) content, which is important for seeing the finer details (Kauffman et al., 2014). A number of neuroimaging studies in human cortical areas of scene and object processing provide support for this coarse-to-fine processing. According to Bullier (2001), faster LSF processing in the dorsal stream guides the slower HSF processing in inferotemporal cortex. Categorisation of visual stimuli may be dominated by LSF or HSF information, depending upon the duration of the stimuli being presented (Schyns & Oliva, 1994). Activation of orbitofrontal cortex (OFC) is elicited by stimuli containing LSF information, 50ms prior to areas in the temporal cortex, but not by HSF-only images, indicating that OFC has an important role in top-down facilitation of image recognition (Bar et al., 2006). LSF modulates HSF processing in broadband images, as measured by EEG in human participants (Petras et al., 2019). When LSF is informative of the image content, HSF contributes to a lesser extent, indicating that coarse information does indeed guide the processing of fine detail.

Developmental progression from lower to higher spatial frequencies appears to be an inherent characteristic of the human visual system. Here we explore whether this progression would alter the representations that emerge in image processing. Interestingly, in the past few years, a new class of models for human visual information processing has become very popular, namely deep convolutional neural networks (CNNs). With the unprecedented success of CNNs on tasks such as image recognition (He et al., 2016; Krizhevsky, et al., 2012; Simonyan & Zisserman, 2015; LeCun et al., 2015 for an overview), these models are now widely used for tasks depending upon visual representations in cognitive and computational neuroscience domains. Jozwik et al. (2017) found that CNNs outperform feature-based models in explaining human similarity judgements. Peterson et al. (2017) demonstrated a high correlation between the activations in fully connected layers of CNNs and human behavioural similarity judgements of animal image pairs. Kubilus et al. (2016) discovered a striking similarity between human behavioural and CNN shape representation. Bracci et al. (2019) found that CNNs classify objects by animacy rather than appearance, and this bias towards animacy is also demonstrated in human judgements and brain representations. Zeman et al. (2020) found that CNNs represent category independently from shape in object images, similarly to human object recognition areas. All together, these studies and more, have demonstrated the strength and breadth of applications in using CNNs as a model of human vision.

Despite these promising results, some studies have shown intriguing, counter-intuitive properties of CNNs that place doubt upon their viability as a model of the human visual system. For example, Jozwik et al. (2017) found that categorical models outperform CNNs in human similarity judgements, concluding that CNNs lack relevant semantic information. Peterson et al. (2017) discovered that major categorical divisions (between animal images) were missing in CNN representations. Multidimensional scaling showed that major categorical divisions were not preserved. Szegedy et al. (2013) observed that minute, imperceptible (for humans) changes to an image can drastically change a CNN’s predictions for that image, which surprisingly, and alarmingly, can generalise to models with different architectures that are even trained under different procedures. Similarly, Nguyen et al. (2015) showed that CNNs can be easily fooled by images generated using an evolutionary (gradient ascent) algorithm. These images, which are unrecognisable to human observers, are categorised by CNNs with a very high confidence level and are referred to as adversarial examples. Given these findings, it is clear that there are further improvements to be made with CNNs, not only in their architectural components and connectivity, but also in the method in which they are trained.

In spite of the convolutional and pooling layers of CNNs being directly inspired by simple and complex cells in visual neuroscience, and that the overall architecture is reminiscent of the LGN-V1-V2-V4-IT hierarchy in the visual ventral pathway (LeCun et al., 2015), the idea of progressively increasing the spatial frequency content of images during training has not yet been implemented. One notable exception was implemented for generative adversarial networks, initially training with low resolution images before continuing with higher resolution images (Karras et al., 2018). This progressive method allowed for these networks to “discover the large-scale structure of images prior to details at increasingly finer scales, as opposed to learning all scales simultaneously” (Karras et al., 2018, p. 2). The training benefits were twofold, by sufficiently decreasing the training time as well as improving the stability in synthesizing both low and high resolution images. While such a training regime demonstrated clear benefits, we note that this study did not include any conjecture or comparison with human vision, and so the question remains as to whether training CNNs under such conditions would bring about better performance changes, or representational changes, that would increase similarity to human observers.

Following on from the culmination of findings from different fields, we hypothesise that training a CNN by simulating the progressive exposure from low to high spatial frequencies would increase the performance to be similar to that shown by human behaviour, as well as increase the similarity between artificial and human visual representations. By encouraging the environmental conditions in which an artificial visual brain would develop to be more similar to that of a biological visual brain, we may potentially overcome some of the current limitations of CNNs as models for human visual functions. In the present study, we extensively analysed the effects of implementing a training protocol for deep neural networks that progressively increases the spatial frequency information of images. We compared our results against a variety of human findings, including behavioural and neuroimaging studies.

## Method

### CNN implementation

For our model architecture, we selected MobileNet (Howard et al., 2017), an efficient, serially-connected CNN with 28 layers. We trained MobileNet on the 2012 ImageNet (ILSVRC) dataset (Russakovsky et al., 2015), which contains approximately 1.2M images from 1000 classes. We implemented three different training regimes (see Table 1):

**Table 1.** Training Regimes of MobileNet

| E | Full training |  |  |  |  |  | Gradual training |  |  |  |  |  | Mixed training |  |  |  |  |  |
| --- | --- | --- | --- | --- | --- | --- | --- | --- | --- | --- | --- | --- | --- | --- | --- | --- | --- | --- |
|  | LR | .1 | .3 | .5 | .8 | UF | LR | .1 | .3 | .5 | .8 | UF | LR | .1 | .3 | .5 | .8 | UF |
| 100 | 0.5 | 0 | 0 | 0 | 0 | 100 | 0.5 | 100 | 0 | 0 | 0 | 0 | 0.5 | 100 | 0 | 0 | 0 | 0 |
| 200 | 0.5 | 0 | 0 | 0 | 0 | 100 | 0.5 | 0 | 100 | 0 | 0 | 0 | 0.5 | 50 | 50 | 0 | 0 | 0 |
| 300 | 0.5 | 0 | 0 | 0 | 0 | 100 | 0.5 | 0 | 0 | 100 | 0 | 0 | 0.5 | 25 | 25 | 50 | 0 | 0 |
| 400 | 0.1 | 0 | 0 | 0 | 0 | 100 | 0.1 | 0 | 0 | 0 | 100 | 0 | 0.1 | 16 | 16 | 16 | 50 | 0 |
| 450 | 0.01 | 0 | 0 | 0 | 0 | 100 | 0.01 | 0 | 0 | 0 | 0 | 100 | 0.01 | 12.5 | 12.5 | 12.5 | 12.5 | 50 |

*Note.* E = epoch, LR = learning rate. Values below the decimal number (cy/deg) and UF (unfiltered images) columns denote the proportion (%) of images at the given stage of training.

1. *Full Training*: We trained the network using unfiltered images, in which no SF information is omitted from the images. This regime is the most commonly used technique and serves as a control condition.
2. *Gradual Training*: We progressively shifted the SF information of images during training. At every 100th epoch, the training set was switched to a set of identical, albeit differently filtered images. There were five different training sets, which included one unfiltered set and four differently filtered training sets, each containing images with higher SF information than the previous (see “Filtering of ImageNet images” section).
3. *Mixed Training*: Taking an approach similar to the Gradual Training method, we changed the training set every 100 epochs, except that a proportion of images from the previous set was retained in the subsequent set (those containing lower SF information).

MobileNet converged in 500 epochs (batch size = 192, steps per epoch = 100). We used stochastic gradient descent (SGD) with Nesterov momentum (0.9) as an optimiser and categorical cross-entropy as a loss function. The initial learning rate was 0.5, which decreased to 0.1 after 400 epochs and decreased again to 0.01 after 450 epochs. We measured validation accuracy using validation images that were also filtered on the same SF levels as the training sets. All training regimes were implemented in TensorFlow.

### Filtering of images

We filtered train and test images in MatLab using lowpass Butterworth filters (Butterworth, 1930) by executing the following procedure. First, images were resized to 224×224×3, which is the training size for MobileNet. Second, to convert the measuring unit of spatial frequency information (cycles per degree, or cy/deg) into an analogous unit used by the Butterworth filters (radius in the frequency domain), we defined the pixels per cycle for given images. We assumed a visual angle of 60 degrees, which is the visual angle of a standard smartphone (iPhone 6). Knowing the dimensions of images (224×224×3) and having a physical measure (60 degrees), we could approximate pixels per degree (224/60). This allowed us to compute spatial frequencies (SF) in cycles per degree (see Equation 1). Radius in the frequency domain was then compared to cycles per degree, providing us with all the required prerequisite information to apply Butterworth filters. Third, we applied four different Butterworth filters to ImageNet, each one with attenuating spatial frequencies higher than a threshold in cycles per degree (cy/deg). The thresholds that we applied, referred to as Spatial Frequency Levels or SFL, were 0.1 cy/deg (SFL1), 0.3 cy/deg (SFL2), 0.5 cy/deg (SFL3) and 0.8 cy/deg (SFL4). The specific values of filters were chosen to mimic the development of infant contrast sensitivity levels found in different studies (see Kiorpes, 2016). The four thresholds represent the approximate peak levels of the CSF at different stages of development, from infancy to eight months, which is considered to be the most intense period in development. Equation (2) defines the Butterworth filter, where *D* is distance from centre (namely 113 pixels, given an image size of 224 × 224), *r* is the radius in the frequency domain and *n* is the order of the filter (all filters were of order 4).

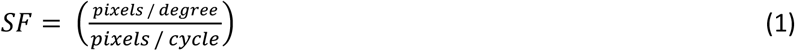

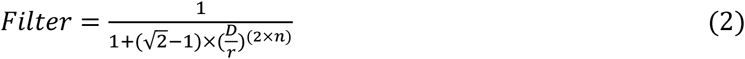

### Representational Similarity Analysis (RSA)

After training the CNNs, we compared representations between them using RSA, which is a framework that allows for quantitative comparisons of internal representations between computational models, and even other modalities such as neural activity and behaviour (Kriegeskorte et al., 2008). For CNN representations, after forward passing the images through the network, we extracted activations from the final fully connected layer and a mid-level convolutional layer (layer 15), in each of the three implementations of MobileNet. From these activations, we constructed representational dissimilarity matrices (RDMs). RDMs represent dissimilarities of activations for each image by measuring correlations (1 – Spearman’s rho). We then compared the RDMs (using Spearman’s rho) to the behavioural and conceptual RDMs from the mentioned studies.

To test for the effect of spatial frequency on representational similarity, we also constructed a set of *hybrid images*, which were inspired by Schyns and Oliva (1994). These images consisted of two different superimposed images, one filtered with a lowpass filter (<0.17cy/deg) and one with a highpass filter (>0.17cy/deg). The set was composed of 18 images. Nine of those images, labelled as HSFΔ, consisted of the same LSF content but different HSF content. The other nine images, labelled as LSFΔ, consisted of the same HSF content but different LSF content. We composed two conceptual RDMs for the set of hybrid images, the so-called LSF and HSF models. We postulated that if an implementation of MobileNet is sensitive only to the HSF content of images, it should represent the images where LSF content is manipulated as more similar than those where HSF content is manipulated (HSF model). In contrast, if an implementation of MobileNet is sensitive only to LSF content of images, it should represent the images where HSF content is manipulated as more similar than those where LSF content is manipulated (LSF model).

To detect any possible changes in representations due to our training protocol, we included two stimulus sets along with the associated CNN, behavioural, and neural similarity data from Bracci et al. (2019) and Bracci et al. (2016; with further analyses by Zeman et al., 2020) for comparison with our trained networks. In both studies, the authors dissociated appearance from the category for each of the stimuli, allowing for a controlled comparison between visual and more conceptual information (see Figures 2 and 3).

**Figure 1.**
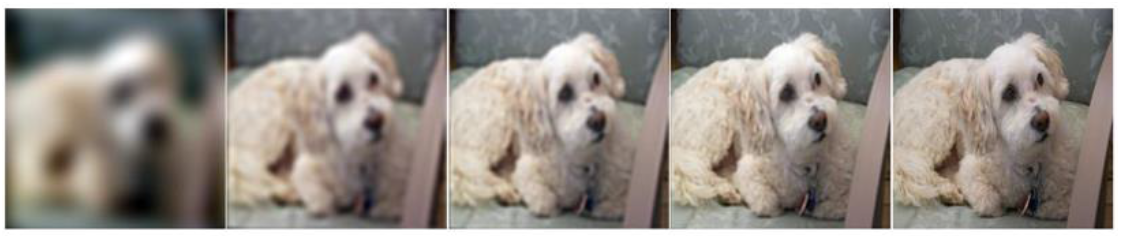
Example of an image from ImageNet with the Butterworth filter applied using increasing SF cut-off thresholds, from left to right: 0.1 cy/deg (SFL 1), 0.3 cy/deg (SFL 2), 0.5 cy/deg (SFL 3), 0.8 cy/deg (SFL 4) and the original unfiltered image.

**Figure 2.**
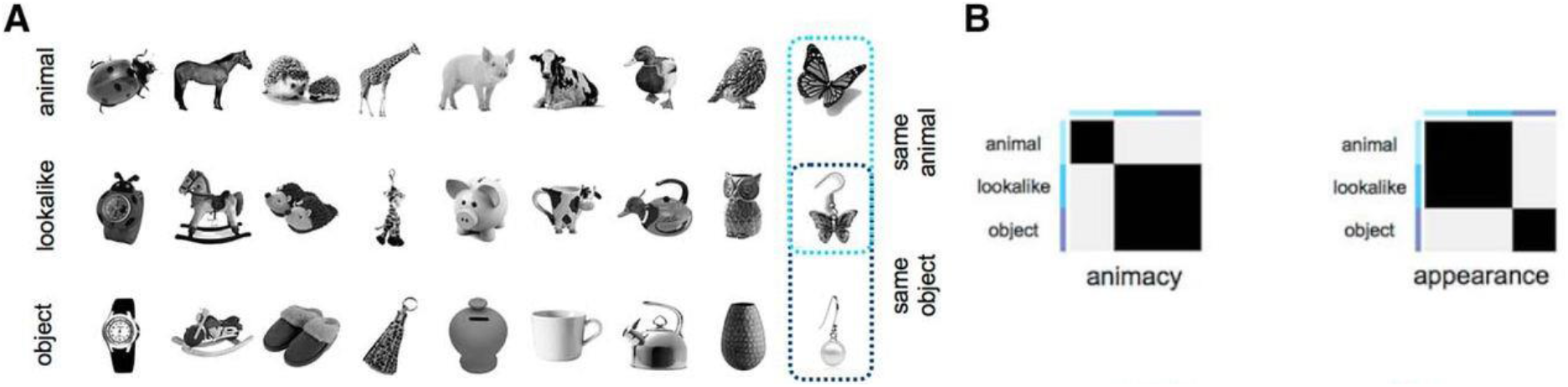
Stimuli (panel A) and conceptual models (panel B) from Bracci et al. (2019). The black areas in the RDMs represent high similarity (1 – Spearman’s rho = 1) and the grey areas represent low similarity (1 – Spearman’s rho = 0). To construct the RDMs, stimuli are numbered left to right, top to bottom row.

**Figure 3.**
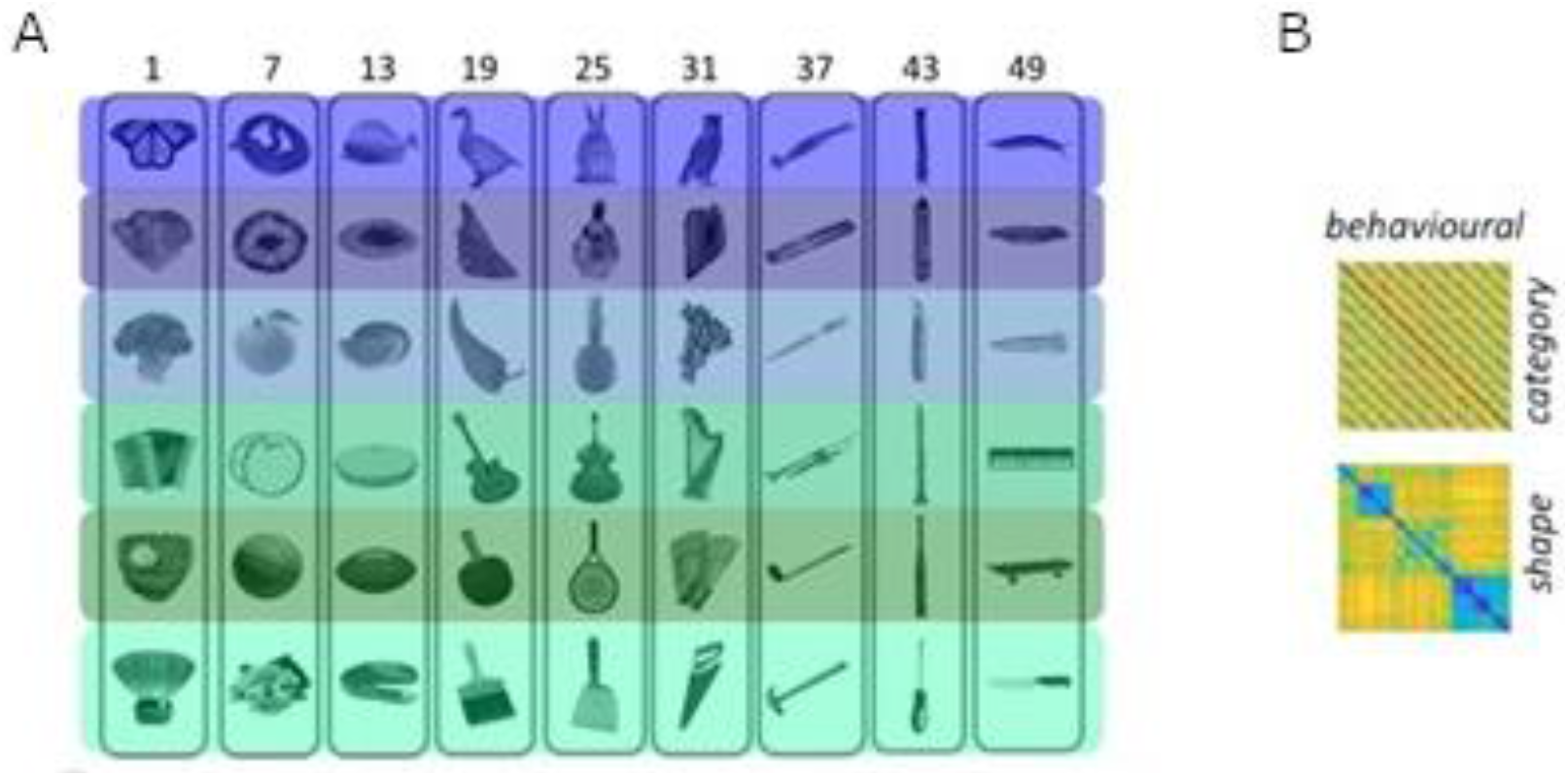
Stimuli (panel A) and behavioural models (panel B) from Zeman et al. (2020). The rows in panel A represent stimuli that share category but differ in shape, whereas the columns represent stimuli that are similar in shape but belong to different categories. The behavioural matrices (panel B) uses blue colour to represent high similarity, whereas yellow represents low similarity.

**Figure 4.**
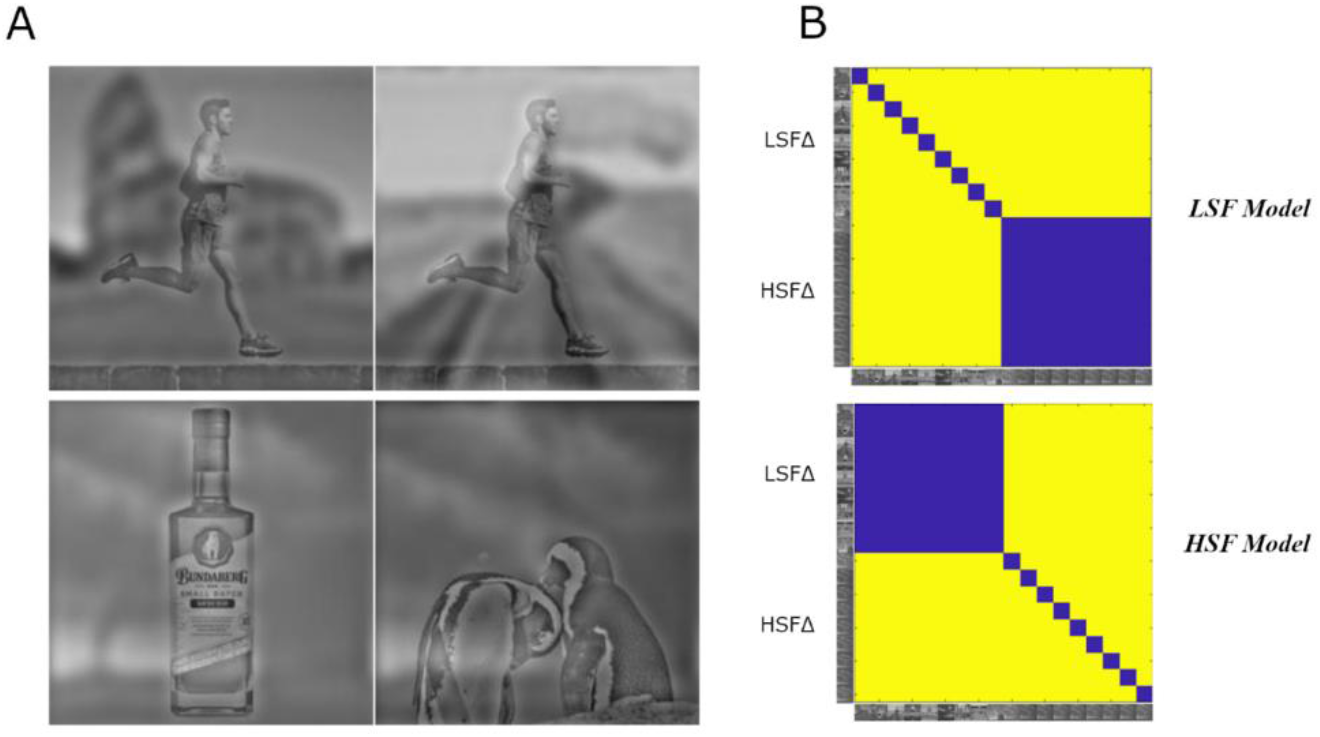
Panel A shows the hybrid stimuli. The top row shows two examples LSFΔ stimuli (left-upper corner), and the bottom row displays two HSFΔ stimuli. Panel B shows our conceptual RDMs. The upper matrix in panel B represents the LSF model. The LSF model shows the greatest similarity between images that contain the same LSF content with varying HSF content, while showing the lowest similarity between images with varying LSF but consistent HSF content. The lower matrix in panel B represents the HSF model. The HSF model shows the greatest similarity between images that contain the same HSF content with varying LSF content, while showing the lowest similarity between images with varying HSF but consistent LSF content.

**Figure 5.**
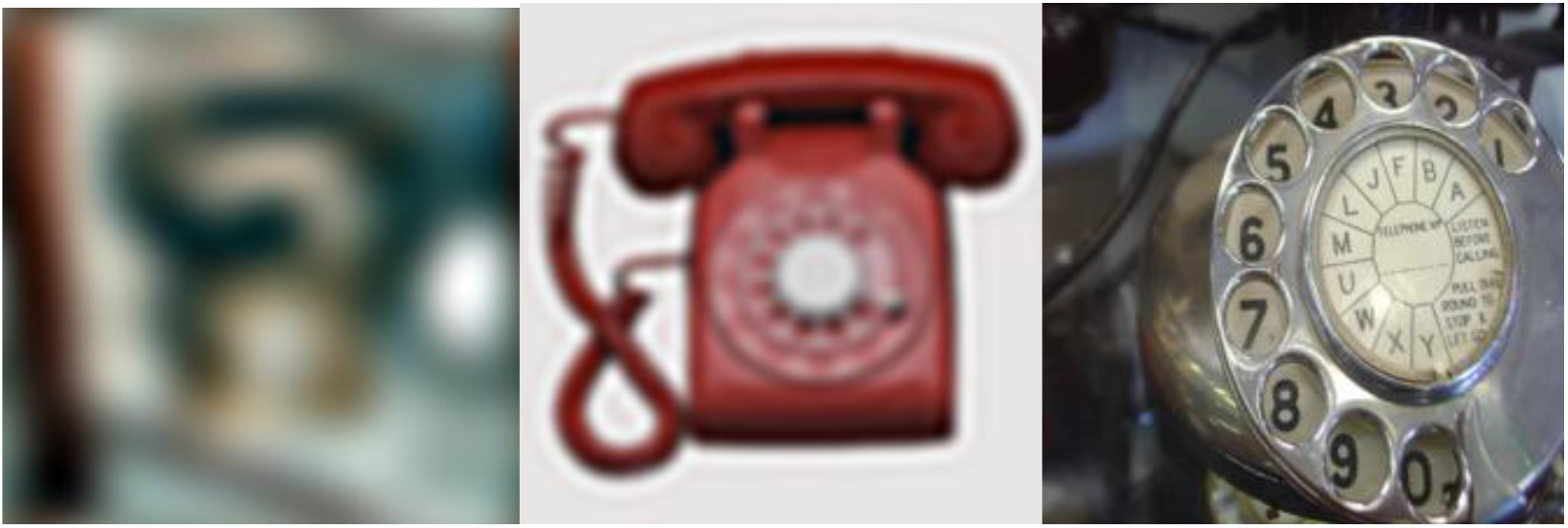
Example of an SFL1, SFL2 and unfiltered image for the category “dial telephone”.

We computed inferential statistics using random permutations and bootstrap methods. To examine if a model RDM correlated significantly with a target (conceptual or behavioural) RDM, we permuted image labels 104 times. We then correlated all the permuted RDMs with the target RDMs. We calculated the p-value by taking the number of correlations with a greater value than the model’s correlation and dividing this by the number of all possible correlations (10^4^). The error bars shown in Figure 7 depict the standard deviations from bootstrapping each model’s correlation 10^4^ times.

### Behavioural experiment

To assess human performance in image classification using differently filtered images, we constructed a behavioural experiment in Psychopy (Peirce et al., 2019). We chose ten categories with consistent performance in the trained CNNs, by extracting activations from the classification layer after feeding in the images from each category and confirming that accuracy levels were within 15% of average performance (15% above or below). These were hammerhead shark (H), cock (C), badger (B), dial telephone (D), planetarium (P), sports car (S), upright piano (U), acorn squash (A), Granny Smith apple (G) and red wine (R). The experiment consisted of two stages. In the first stage, the “training phase”, participants viewed 10 unfiltered examples of each category in a randomised order. An image was displayed indefinitely, and participants were required to press the letter of the corresponding category in order to continue. The “testing phase” followed. In this phase, participants were presented with images in a randomised order from the 10 categories with 3 spatial frequency cut-offs: 0.1 cy/deg (SFL 1), 0.3 cy/deg (SFL 2) and no cut-off (unfiltered). Each image was presented for 150ms. After the elapsed time period, the participant was required to select the letter corresponding to the image (e.g. a for acorn squash). Participants classified 30 images per category, with 10 per spatial frequency level (for a total of 300 images). All images and exemplars were only shown once. To compare human performance in the 10-way categorisation task with CNN performance, we extracted the activations from the classification layer only for the 10 categories. Categorisation was correct if the activation of the corresponding category had the highest value over the other 9 possible outcomes (referred to as “top-1 accuracy”, in this case for a choice among 10 categories instead of the 1000 categories in ImageNet).

The experiment included 28 participants (11 males, 15 females and 2 “other”), with an average age of 23.54 years (*SD* = 3.13), who participated through the online platform Pavlovia.com. The experiment included an agreement to ensure informed consent prior to testing. Procedures were approved by the KU Leuven Social and Societal Ethics Committee.

## Results

### Training and validation accuracy with the three training regimes (Figure 6)

The MobileNet trained with the Full training converged to the highest training accuracy (74%) and validation accuracy (60%) for unfiltered images. However, it performed very poorly (0%) on the lowest spatial frequency level SFL 1 (0.1 cy/deg). Accuracy increased with images containing higher spatial frequency information (21% for SFL 2, 36% for SFL 3 and 48% for SFL 4). With Gradual training, MobileNet showed increasingly better performance during learning for the first 100 epochs, during which it was only exposed to low SF images (SFL 1). However, there was an immediate drop (to 0%) in performance after switching to the image set containing higher SF information. Simultaneously, there was a very rapid stepwise increase of accuracy, from 0 to around 50%, for the sets containing higher SF information. With Mixed training, we again observed a very rapid increase in performance of MobileNet after the 100th epoch for images containing higher SF information, yet there was no drop in performance for images containing the lowest spatial frequency information, which instead converged further. Interestingly, the Mixed training policy was the only training regime that allowed the model to reach a sufficient level of performance on the images containing only the lowest SF information (32%), while maintaining relatively high levels of accuracy on the images with higher cut-off levels and unfiltered images (48% for SFL 2, 50% for SFL 3, 51% for SFL 4, 60% for training accuracy and 51% for validation accuracy on unfiltered images).

**Figure 6.**
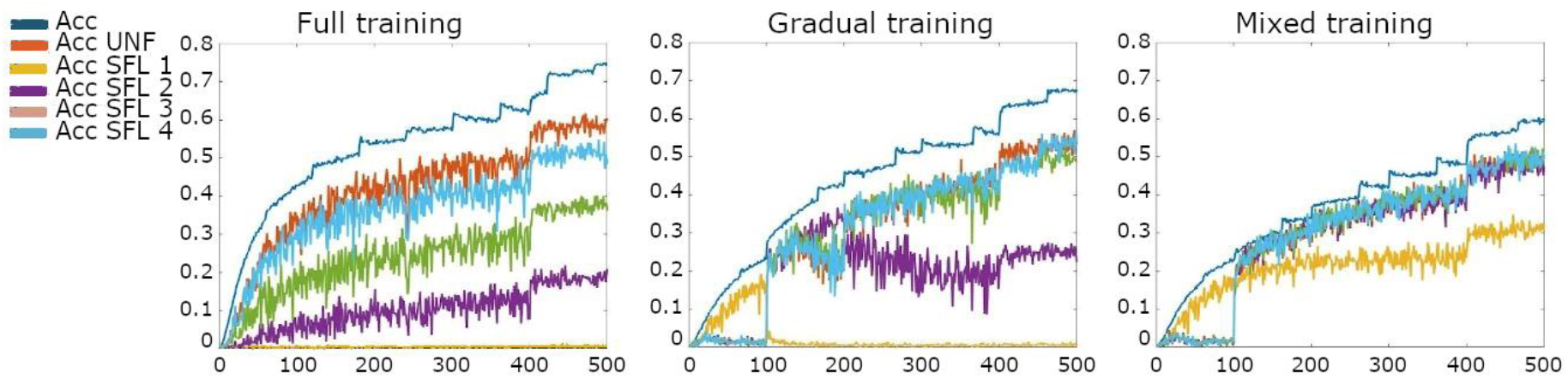
Top-1 performance of MobileNet trained with the Full, Gradual and Mixed (from left to right) regimes on images with different spatial frequency levels (SFL 1 = 0.1cy/deg, SFL 2 = 0.3 cy/deg, SFL 3 = 0.5 cy/deg, SFL 4 = 0.8 cy/deg, Acc UNF = validation accuracy on unfiltered images, Acc = training accuracy).

### Comparison with human performance for low-pass filtered images

We tested how well human participants would classify low-pass filtered images, with results shown in Figure 7. Note that participants did not receive training with such images in the context of the experiment. As the behavioural experiment included only ten categories, the performance of the CNNs was also calculated for the same ten category task. If we consider the most strongly filtered images as SFL 1, the Full training policy, which was never presented with these filtered images during training, performed very poorly. Although the Gradual training was initially trained on filtered images, its performance on SFL 1 did not exceed that of the Full training policy. Both human participants and the Mixed training model reached relatively high accuracy with the SFL 1 images, the latter surpassing human performance by about 10%. With the SFL 2 condition, the Full training shows a marked drop in performance compared to unfiltered images, which was not present in the other models and in human performance.

**Figure 7.**
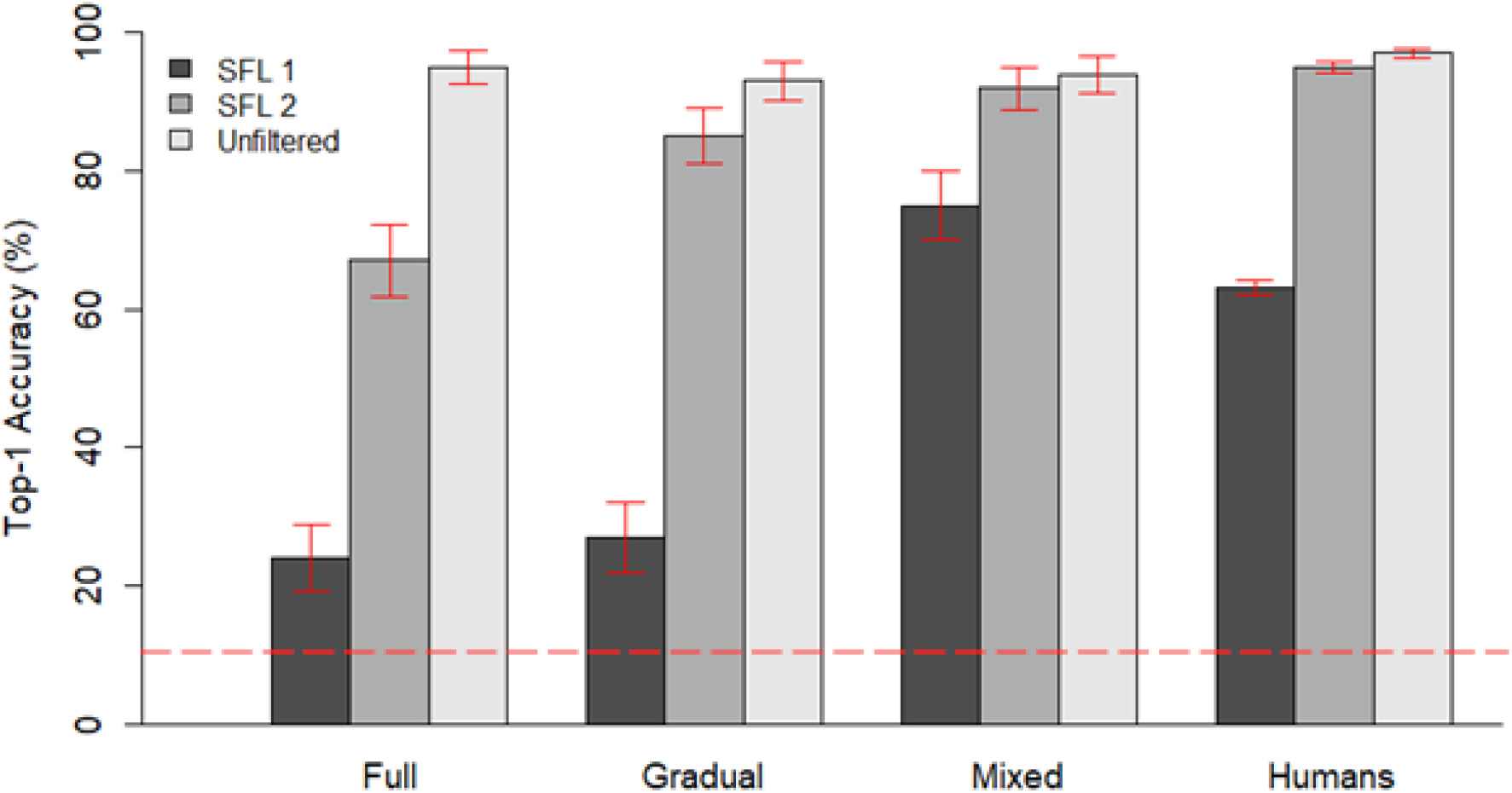
Performance of MobileNet trained with different regimes and humans on a 10-way categorisation task containing images with different spatial frequency information (Dark grey: SFL 1 = 0.1 cy/deg, mid-grey: SFL 2 = 0.3 cy/deg, light grey: Unfiltered). In the case of MobileNet, error bars indicate the binomial confidence interval, whereas with humans, they indicate standard error of the mean (*SEM*). The horizontal dashed line indicates the chance level (10%).

Overall, it seems that an extensive and continuous exposure to filtered images during training is necessary to allow deep learning models to reach the capacity of humans to recognise images that only contain low spatial frequency content.

### RSA with hybrid stimuli

We investigated whether the training affected the representational similarity of the hybrid images that combine different information at low and high spatial frequencies. The results are shown in Figure 8. Both layers show similar effects of the type of training. This suggests that the differentiability that the networks learn to distinguish images with low SF content is present in earlier layers of the network, and not only in the final classification layer.

**Figure 8.**
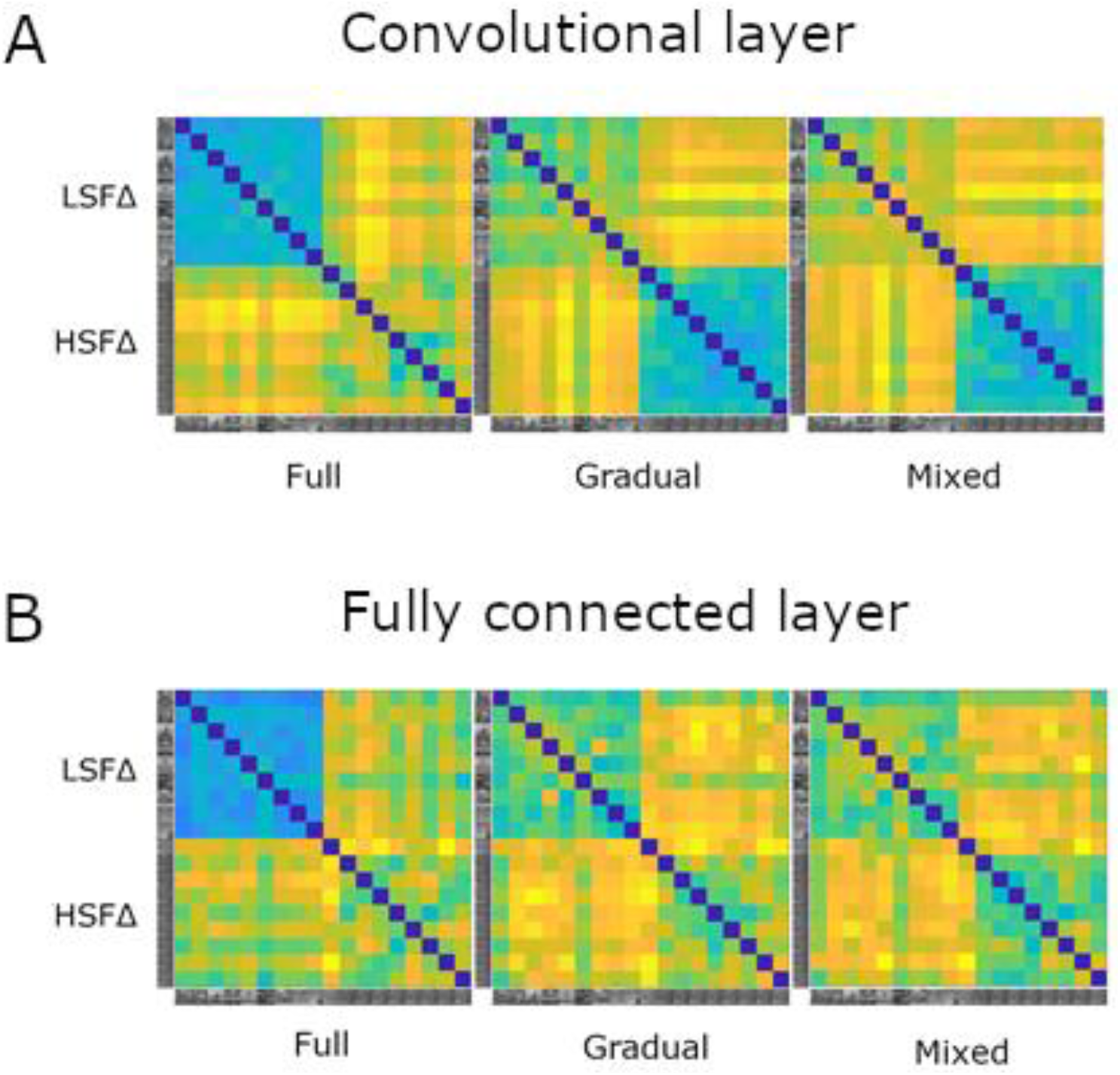
From left to right: RDMs for the Full, Gradual and Mixed training of MobileNet for the mid-level (layer 15) convolutional layer (panel A) and the fully connected layer (panel B). Blue denotes high similarity, whereas yellow denotes low similarity. Labels on the left side show how the stimuli are ordered.

Figures 9–10 display the correlations of these CNN representations with the conceptual LSF and HSF model. In the fully connected layer, only the model trained with Mixed training regime reached significant positive correlation for both the LSF and HSF model. In other words, the Mixed training model was sensitive to the manipulation of both the low and high spatial frequency content of the images. In contrast, the Full training model had a significantly negative correlation with the LSF model and the highest positive correlation with the HSF model. This indicates that after Full training, the network is capable of differentiating images with varied HSF content, but not images with varied LSF content, which elicit similar activations. Findings were similar between the convolutional layer and the fully connected layer, with the exception that in Gradual training, the convolutional layer correlated with the LSF model. Overall, a Mixed training regime is required for a model to become sensitive to both the low and high spatial frequency information in images.

**Figure 9.**
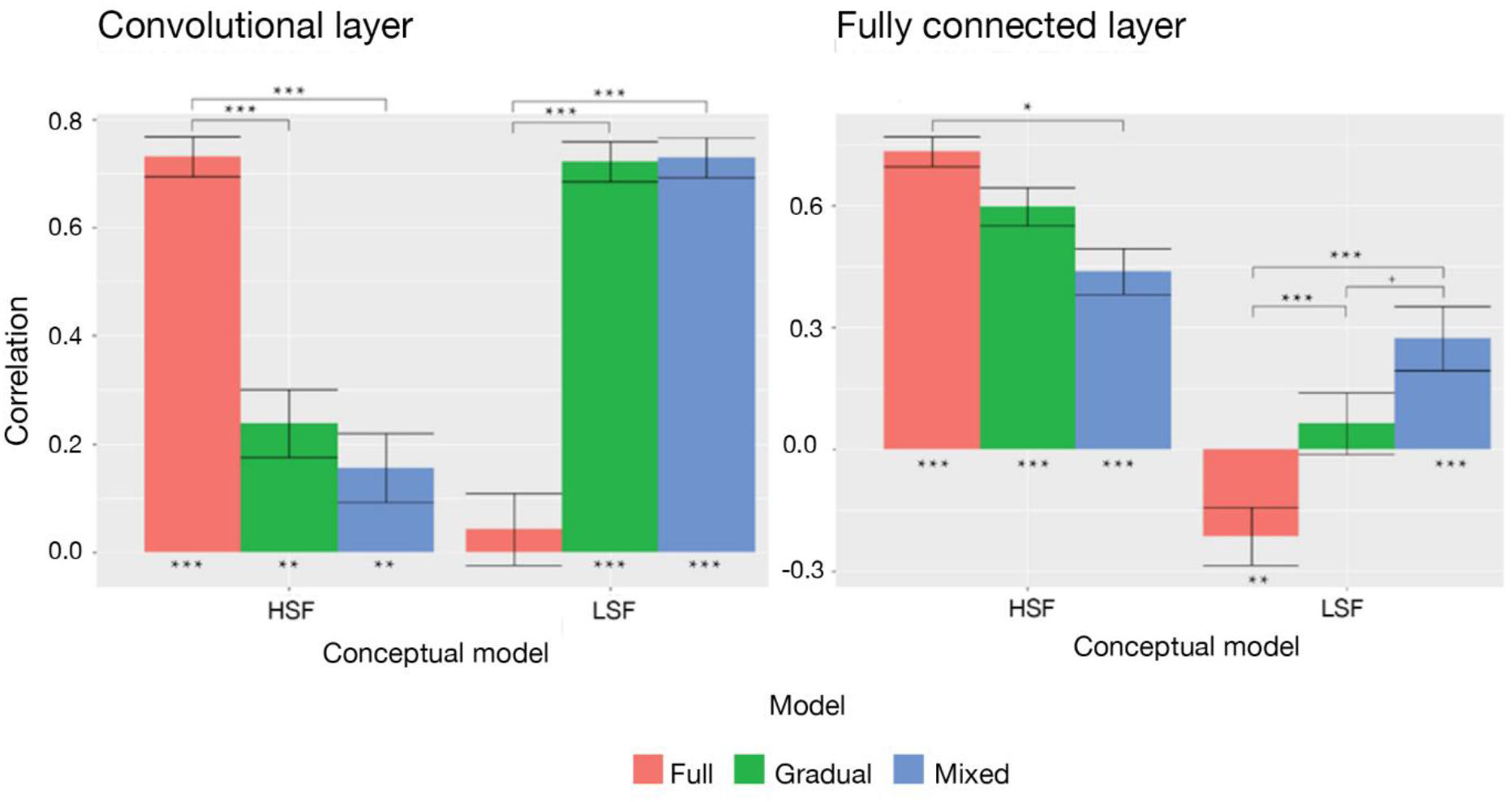
Left: Correlations between conceptual models for hybrid stimuli and the convolutional layer (layer no. 15) of different MobileNet models (red: Full training, green: Gradual training, blue: Mixed training). Right: Correlations between conceptual models for hybrid stimuli and the fully connected layer of different MobileNet models. Standard errors represent the standard deviations of 10^4^ bootstraps. *** = p < 0.001, ** = p < 0.01, * = p < 0.05, + = p < 0.10

**Figure 10.**
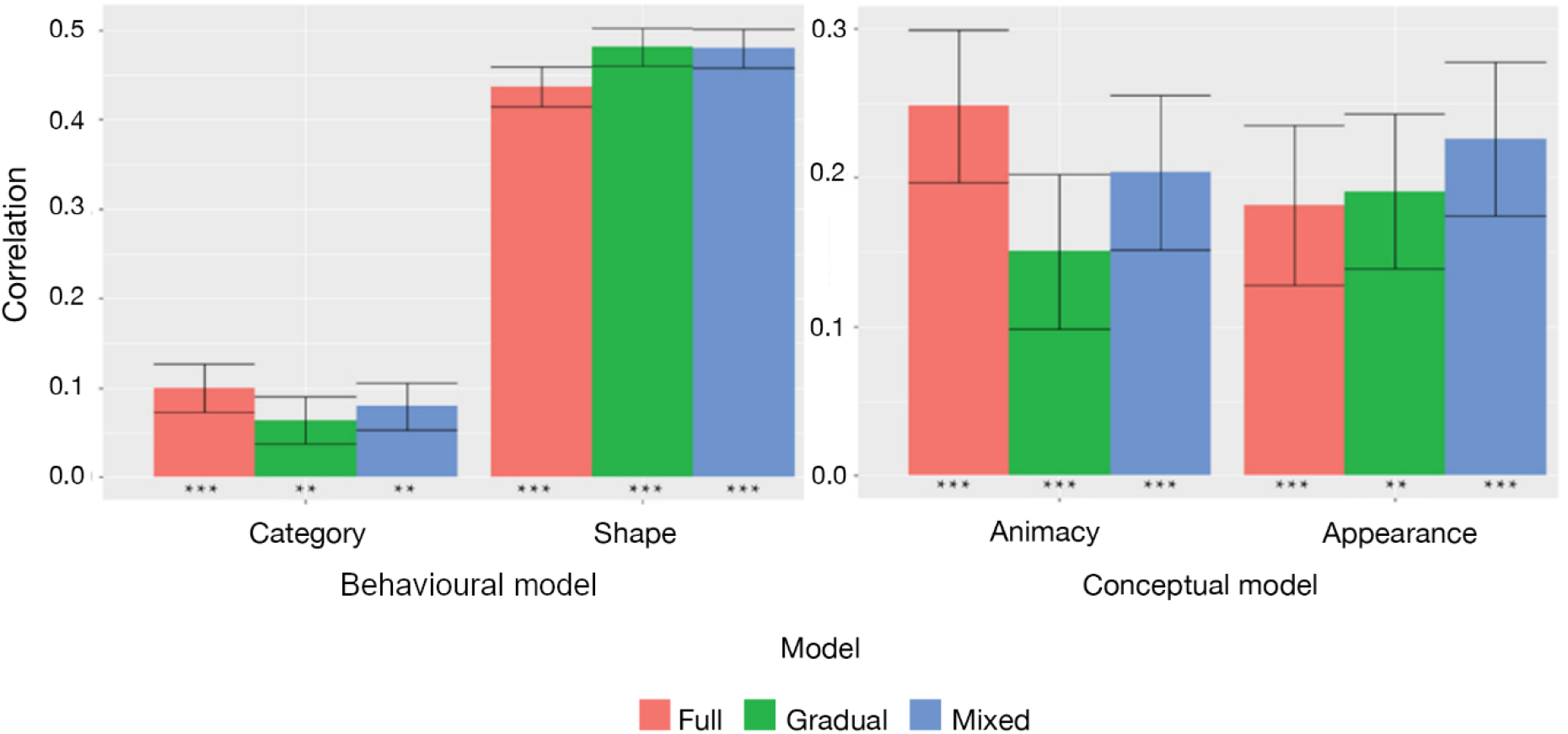
Left: Behavioural model (from Zeman et al., 2020) correlations with the fully connected layer of different MobileNet models. Right: Conceptual model (Bracci et al., 2019) correlations with the fully connected layer of different MobileNet models. Standard error bars represent the standard deviation of 104 bootstraps. *** = p < 0.001, ** = p < 0.01, * = p < 0.05, + = p < 0.10

### RSA with stimuli from Bracci & Op de Beeck (2016) and Bracci et al. (2019)

By now we have shown that a Mixed Training regime is important to obtain a CNN that is able to recognise objects from low-spatial frequency content and take this content into account in object representations. Next, we investigated whether such a training regime would also affect the representational similarity for stimulus sets that do not include an explicit manipulation of spatial frequencies.

For the stimulus set of Bracci & Op de Beeck (2016), all three training regimes of MobileNet preferred shape over category. In addition, they also showed a smaller, yet significant correlation with the category-related behavioural similarities, as it was also shown before for other networks trained with unfiltered images by Zeman et al. (2020). There are no meaningful effects of the training regime, so the extent and timing of training with low-pass filtered images does not affect the presence of a multi-feature representation in a CNN.

Bracci et al. (2019) showed that CNNs pre-trained on ImageNet (VGG-19 and GoogleNet) have a strong bias for the Animacy model over the Appearance model, which puts CNN models at odds with human perception and neural responses. We also see a stronger correlation for the Animacy model with MobileNet when trained with unfiltered images. Nevertheless, the bias towards Animacy in the late fully connected layers was much larger in Bracci et al. (2019).

The bias towards Animacy is no longer present when training includes low-pass filtered images. Given the variability (shown by large error bars), we cannot state that this effect is significant, so the findings are inconclusive about a meaningful effect of training regime upon the nature of representations investigated with this stimulus set.

## Discussion

In our study, we thoroughly examined the effect of a training regime with progressive exposure to images with increasingly finer spatial frequencies, akin to biological vision. This training method allowed us to determine the effect on deep neural network performance and representations and investigate whether this bears resemblance to human performance and representations.

The most widely used training regime in computer vision and neuroscience communities, which exposes a model only to unfiltered images, is not able to accurately classify visual stimuli that contain only low spatial frequency information. In both the 1000-way and the 10-way task (Figures 6 and 7), the Full training model demonstrated very low levels of categorisation accuracy when presented with images that had a cut-off spatial frequency of 0.1 cy/deg (SFL 1). Comparing this outcome to human performance, participants demonstrated much greater accuracy with such stimuli, surpassing the performance of the Full training model by almost a factor of 3, despite most of the spatial frequency information being omitted (SFL 1). Notably, even with initial exposure to SFL 1 images, the model’s performance on these images was not sustained unless a proportion of those images were retained in the training set. This was visible in the performance of the Gradual training model (Figure 6), with an immediate drop after switching to images with a higher cut-off (SFL 2). We observed a similar, albeit less dramatic fall in performance, for SFL 2 images at epoch 200. There was no apparent drop in accuracy for higher cut-off levels (SFL 3 and 4).

Our findings with Gradual training are peculiar. Given the cumulative nature of spatial frequency information, where each set of images with a higher SF cut-off contains information from the lower SF cut-off plus new, higher SF information, we would expect that initial exposure is enough to calibrate the weights of a model in a way that would later provide sensitivity to low spatial frequencies. However, this was not the case. Apparently, providing images with lower levels of spatial frequency information is necessary throughout the whole training process to sustain high levels of accuracy. This could be a result of the non-additive way in which weights are computed in the models. From this perspective, the relation of non-linear functions of the model to those of the human visual system is put into question. Although CNNs reach high levels of accuracy in image categorisation tasks, they might achieve this in an “inherently” different way to humans. To express it differently, the ability to achieve similar results, in terms of outstanding categorisation accuracy, might stem from different underlying processes.

The possibility of divergence between the underlying processes of the human visual system and of CNNs is further supported by the findings with the Full training. Even for the SFL 2 images, the accuracy of the Full training model was substantially lower than human participants, whose accuracy is almost equivalent to the unfiltered images (Figure 7). Indeed, if we look at an example of an SFL 2 image (Figure 1), the changes caused by the filter are so minute that they are barely registered by the human eye. Yet the Full training model struggled with such images. These limitations were solved using a simple technique: retaining the lower SF images in the training set the whole time, as we did in the Mixed training regime.

The implication that the Full training model was not sensitive to LSF content was reaffirmed by results from RSA. We can see that the Full training model was sensitive to manipulation of HSF content, but not LSF content, as it correlates significantly only with the HSF model. On the other hand, the Mixed training was sensitive to both manipulations, which is reflected in significant correlations with both the LSF and HSF model (Figure 9). We must acknowledge that there is possibly a trade-off between sensitivities at different SF levels. The enhanced sensitivity for LSF information results in deteriorated sensitivity for HSF information. Thus, the Full training model surpasses the Mixed training model in terms of classification accuracy for unfiltered images in the 1000-way task, and in correlation levels with the HSF model, where we can see an almost linear decrease in the Full-Gradual-Mixed order (Figure 9). Nevertheless, this trade-off does not have such a large magnitude, and the Mixed Training model produces very sensible behaviour, incorporating both LSF and HSF information.

Furthermore, we examined if different training regimes affected internal CNN representations of visual stimuli, where spatial frequencies were not directly manipulated (Figure 10). Correlations with RDMs from Zeman et al. (2020) did not show any significant changes. All three models correlated significantly with both behavioural models but preferred shape over category. No statistically significant differences between models were found. Likewise, the results for Bracci et al. (2019) stimuli showed significant correlation for both conceptual models. There are some apparent fluctuations across training regimes but given the standard errors, we cannot conclude that there are significant differences. The findings with the Full training model tend to be most similar to the findings reported by Bracci et al. (2019), who demonstrated a much higher preference for the Animacy model than for the Appearance model. We can still see a slight preference for Animacy with the Full training model, whereas this preference disappeared in the Mixed training model. Further tests with larger stimulus sets will be needed to investigate whether such effects would be statistically robust.

To sum up, we showed that CNNs trained in the usual way display some properties that make their similarity to human representations questionable. Firestone (2020) argued that when comparing human to CNN behaviour, we should make a distinction between competence and performance, this distinction being a methodological tool to compare the behaviour of different organisms (infants-adults, animals-humans, etc.). Competence relates to the underlying capabilities of an organism, whereas performance relates to behavioural output, which does not necessarily reflect the full capacity of organisms’ underlying resources. Such an approach can facilitate differentiation between organisms at superficial and deep levels. Differences in behaviour can be the result of a variety of factors. These include human constraints, machine constraints and non-aligned species-specific tasks. One reason that the Full training regime performed so badly on LSF images might have been a limitation from the digital input resolution. Humans view images on displays, using their lens, which can further distort the resolution (Firestone, 2020). In our case, the human constraint – limited visual acuity – could play a vital role in incorporating LSF information for object categorisation. In fact, modelling a human fovea (Deza & Konkle, 2020) or primary visual cortex (Dapello et al., 2020) at the front of CNNs can increase their robustness to adversarial examples. Note that adversarial examples usually include subtle changes to images at high spatial frequencies. The human burden of limited visual acuity makes these manipulations imperceivable and thus does not affect human behaviour (Firestone, 2020). It would be interesting to examine whether the addition of a foveal model or primary visual cortex would improve CNNs’ capability to use low spatial frequency information.

Similarly, we can ask whether our Mixed training regime would improve CNNs’ resistance to adversarial examples. As already mentioned, like humans, the Mixed training was capable of using LSF information, and thus showed much higher accuracy for such images than the Full training regime, which was trained by the method that is normally used in computational neuroscience. This technique was done by keeping a proportion of images with a lower cut-off frequency filter throughout the whole training. Such a method restrained the model from only focusing on high spatial frequencies. In this way we induced a “human-like” constraint, similarly to what Deza and Konkle (2020) have done with adding a model of the human fovea. Despite the similarity in performance, the question of concordance between the underlying processes remains. To elaborate on this point: Bar et al. (2006) demonstrated that LSF information affected the processing of HSF information in a top-down manner. None of our models were made to simulate top-down processes, which require recurrent connections. An intriguing pathway of research would be to implement our method of Mixed training on a Recurrent Neural Network. A further concern is the problem of generalisability. Our training protocols were implemented on a specific CNN (Mobilenet) with a specific architecture and other properties. However, given that all CNN networks are very similar in how image content is treated in the convolutional layers, we expect that the major benefit of a Mixed training would generalise widely across networks, with possibly some variation in the effect size of this benefit.

Our findings provide some direct implications for real-world applications, particularly in the field of autonomous driving with incorporated real-time processing of camera images. According to Zang et al. (2019) one of the critical issues of autonomous driving systems is their performance under adverse weather conditions, such as rain, snow and fog. These conditions distort the pixel intensities and thus lower the quality of images. For example, raindrops can create patterns that blur the edges in a scene and, in doing so, impede the recognition of objects (Kurihata et al., 2005). Of special interest is the perception of scenes under foggy conditions. Whereas scenes are composed of a broad spectrum of spatial frequencies under normal viewing conditions, the frequency components are concentrated at low spatial frequencies under foggy conditions (Zang et al., 2019). Thus fog acts like a lowpass filter that blurs the finer details of an image. As we have shown, such a filter can have detrimental effects for deep neural networks that had not been exposed to filtered images. The success of our Mixed training protocol suggests the possibility of its implementation in autonomous driving systems for improved performance.

## Acknowledgments

We would like to express our gratitude to Anne-Sofie Maerten for her helpful technical insights and improvements towards our use of deep neural networks.

